# Speed-related but not detrended gait variability increases with more sensitive self-paced treadmill controllers at multiple slopes

**DOI:** 10.1101/2021.01.27.428459

**Authors:** Cesar R. Castano, Helen J. Huang

**Affiliations:** Department of Mechanical and Aerospace Engineering, University of Central Florida, Orlando, FL, United States; Disability, Aging, and Technology Cluster, University of Central Florida, Orlando, FL, United States

## Abstract

Self-paced treadmills are being used more frequently to study humans walking with their self-selected gaits on a range of slopes. There are multiple options to purchase a treadmill with a built-in controller, or implement a custom written self-paced controller, which raises questions about how self-paced controller affect treadmill speed and gait biomechanics on multiple slopes. This study investigated how different self-paced treadmill controller sensitivities affected gait parameters and variability on decline, level, and incline slopes. We hypothesized that increasing self-paced controller sensitivity would increase gait variability on each slope. We also hypothesized that detrended variability could help mitigate differences in variability that arise from differences in speed fluctuations created by the self-paced controllers. Ten young adults walked on a self-paced treadmill using three controller sensitivities (low, medium, and high) and fixed speeds at three slopes (decline, −10°; level, 0°; incline, +10°). Within each slope, average walking speeds and spatiotemporal gait parameters were similar regardless of self-paced controller sensitivity. With higher controller sensitivities on each slope, speed fluctuations, speed variance, and step length variance increased whereas step frequency variance and step width variance were unaffected. Detrended variance was not affected by controller sensitivity suggesting that detrending variability helps mitigate differences associated with treadmill speed fluctuations. Speed-trend step length variances, however, increased with more sensitive controllers. Further, detrended step length variances were similar for self-paced and fixed speed walking, whereas self-paced walking included substantial speed-trend step length variance not present in fixed speed walking. In addition, regardless of the self-paced controller, subjects walked fastest on the level slope with the longest steps, narrowest steps, and least variance. Overall, our findings suggest that separating gait variability into speed-trend and detrended variability could be beneficial for interpreting gait variability among multiple self-paced treadmill studies and when comparing self-paced walking with fixed speed walking.

## Introduction

Self-paced treadmills (also known as user-driven, active, or adaptive speed treadmills) are typically motorized treadmills that modulate belt speeds to match the subject’s walking speed and are becoming more prevalent in gait laboratories. Self-paced control algorithms can use ground reaction forces, marker locations, and gait parameters, among other real-time measures, to determine when and how much to speed up or slow down the treadmill belts. Often, the intent for using self-paced treadmills is to match overground walking speeds and allow for less constrained gait [1–3]. As such, subjects can walk at their self-selected walking speed for several minutes and hundreds of strides while preserving natural gait fluctuations on self-paced treadmills [4–6]. Self-paced treadmills can also be set at incline and decline slopes to study self-selected uphill and downhill gaits [7], overcoming limitations of overground sloped walking studies that are often constrained to short (10-20 meter) ramps or available outdoor terrains [8–11].

Several studies have investigated differences between self-paced and fixed speed walking on a level treadmill. Unlike self-paced walking where the treadmill adjusts to the subject’s gait, the subject adjusts their gait to match the constant treadmill speed during fixed speed walking. Studies showed that stride variabilities, muscle activities, and walking speed during self-paced treadmill walking were more similar to overground walking than fixed speed treadmill walking [2,6,12]. Peak anterior ground reaction forces also increased on a user-driven treadmill compared to a fixed speed treadmill [2]. On an active (self-paced) treadmill, the sensorimotor cortex was more engaged compared to passively walking on the (fixed speed) treadmill [13]. Additionally, self-paced treadmills have helped demonstrate that exoskeletons, visual feedback, and mechanical perturbations can shift a subject’s preferred walking speed [14–16].

As more groups incorporate self-paced walking into their studies, more customized self-paced controllers are being offered as a built-in mode (ex. Motek Medical’s M-Gait, used in this study) or developed and implemented on treadmills already in a laboratory or on new treadmills that lack a self-paced mode option [1,2,17–21]. A previous study evaluated step kinematics and step variability for three control algorithms when walking on a self-paced treadmill at a level slope resulted in a range of speed fluctuations that resulted in similar step kinematics but a wide range of significantly different step length standard deviations (0.037-0.062 m) [6]. These differences in step length variability may make comparing and interpreting findings across multiple self-paced walking studies difficult. Identifying spatiotemporal gait metrics that can account for differences in speed fluctuations and treadmill responsiveness of self-paced treadmill controllers may improve our understanding of self-paced walking.

Gait parameters, particularly those with a strong relationship with speed such as step length [22], are likely to be sensitive to differences in self-selected gaits and speed fluctuations. Gait parameters have both long-range correlations over hundreds of steps [23] and short-range dependencies on preceding strides [24], which may modulate walking speed over short distances [25]. By removing the speed relationship from step length during overground walking, step length variability can be separated into speed-related (speed-trend) and speed-independent (detrended) components. These components may represent long-term and short-term active balance control [26]. A few other overground gait studies also removed speed-related trends to analyze detrended gait variability to avoid speed-related effects [27,28]. If self-paced treadmill walking is similar to overground walking, then detrended gait variability may be beneficial for analyzing self-paced treadmill gait. Recently, two studies reported conflicting results regarding the reliability of spatiotemporal variability during self-paced walking using two different self-paced treadmill systems. One study showed that spatiotemporal gait variability on a self-paced treadmill is reliable for within-session and between sessions [29], whereas another study found significant differences in spatiotemporal variability between sessions [30]. One potential factor that could affect reliability is differences in speed dynamics due to the self-paced treadmill systems.

The primary purpose of this study was to investigate how self-paced treadmill controller sensitivity affects spatiotemporal gait parameters on different slopes (decline, level, and incline) and to determine whether detrended gait variability may mitigate differences in treadmill speed fluctuations of self-paced controllers. We hypothesized that increasing self-paced controller sensitivity would increase speed fluctuations and gait variability within each slope. We also hypothesized that detrended gait variability could help mitigate the expected differences in speed fluctuations and would be similar across sensitivities within a slope. Because studies often use a single sensitivity (or set of controller parameters) for all conditions, a secondary purpose was to investigate the effect of slopes on spatiotemporal gait parameters with each self-paced controller sensitivity. We hypothesized that regardless of sensitivity, self-paced walking speed would be fastest and gait variability would be smallest on the level slope compared to decline and incline slopes for each sensitivity.

## Materials and Methods

Ten healthy young adults (22.6±3.5 years; 4 females) with no musculoskeletal or neurological conditions participated in this study and provided informed written consent. The University of Central Florida Institutional Review Board approved the protocol and consent form.

Subjects walked on a self-paced instrumented treadmill (M-Gait System and D-Flow software version 3.28, Motek Medical B.V., Amsterdam, Netherlands) as motion capture data using the 16-marker OptiTrack “Conventional Lower Body” marker set was recorded (22 cameras, Prime 13 and 13W, OptiTrack NaturalPoint Inc., Corvallis, Oregon). The self-paced controller was designed by the manufacturer and allows users to adjust 4 parameters in the D-flow graphical user interface. These parameters set 1) whether to set the baseline position to be the center of the treadmill or the person’s initial position when the controller is engaged, 2) whether to use the “new algorithm,” 3) whether to stop the self-pace controller when the person stops taking steps, and 4) the gain or sensitivity of the controller. We used the center of the treadmill as the baseline position, which also coincides with the origin of lab’s coordinate system, the new algorithm, the option to stop the self-paced controller when the person stops, and different sensitivities (0.5, low; 1.0, medium; 1.5, high). The new algorithm was released in D-flow (v. 3.10) to compensate better for large accelerations or decelerations and is less sensitive to the natural pelvic movements during walking. To visualize the differences between the new algorithm used in this study and the old algorithm reported in Sloot et al. and used in several studies [4,7], we provide plots of the belt speed, center of mass position, and center of mass velocity in the appendix. Conceptually, the algorithm uses the relative position of the subject’s approximate center-of-mass (average of 4 pelvis markers’ positions) with respect to the center of the treadmill to make a “correction” that adjusts the treadmill belt speeds (Fig. 1). The sensitivity parameter controls how rapidly the controller corrects the treadmill belt speeds to adjust the subject’s position on the treadmill. The equation for the correction used in the old algorithm was reported in Sloot et al., but the specific equation of the “new algorithm” is not disclosed.

**Figure 1:**
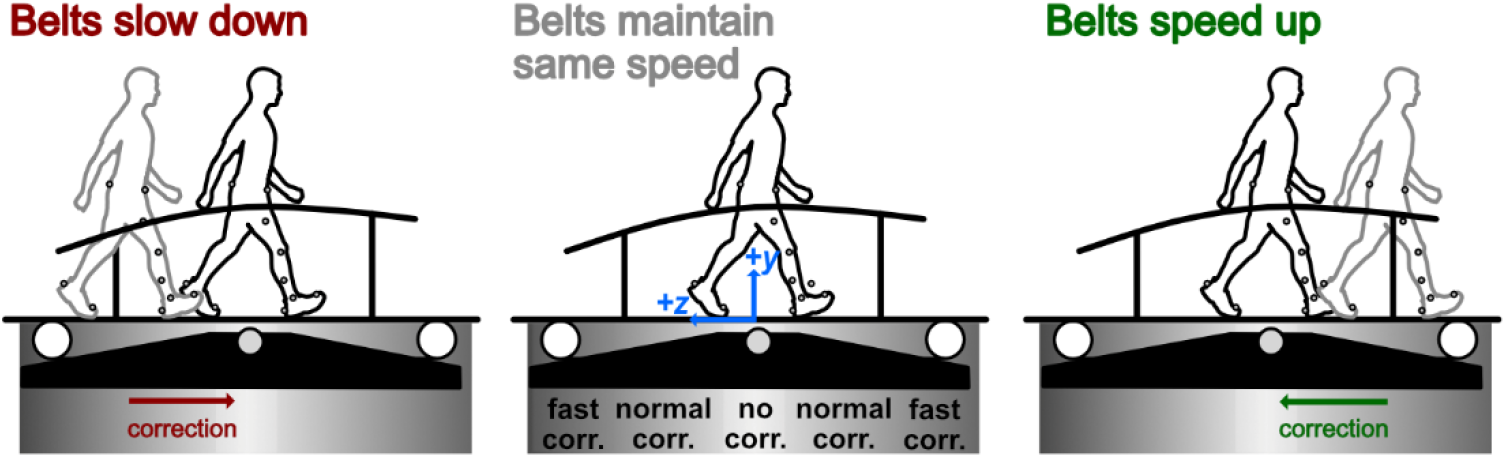
Schematic illustrating the self-paced controller concept. When the subject moves in front of the center point of the treadmill, the treadmill belts speed up (green correction arrow). When the subject moves behind the center point, the treadmill belts slow down (red correction arrow). When the subject remains near the center point, the treadmill belts do not change speed.

### Experimental Protocol

Subjects first walked on the treadmill with a range of fixed speeds and self-paced controller sensitivities during a brief familiarization period. They also completed a 10-meter walk test to identify their level overground walking speed. Subjects then completed nine 5-minute self-paced treadmill walking conditions, which were the combinations of controller sensitivities (low, medium, high) and slopes (decline, −10°; level, 0°; incline, +10°) in a randomized order with brief 1-2 minute rest periods between conditions. Subjects then completed 3 subject-specific fixed speed conditions, one for each slope. The subject-specific fixed speeds were calculated as the average walking speed (using from the last 2.5 minutes of a condition) of the 3 controller sensitivities on a slope for each subject. While all subjects performed the fixed speed conditions, a file naming error resulted in just five subjects having complete fixed speed data sets.

### Spatiotemporal Gait Parameters

Using custom MATLAB (Mathworks, Inc., Natick, Massachusetts) scripts, we first resampled the treadmill belt speed data from 333 Hz to 240Hz to match the motion capture system. We applied a low-pass filter (zero-lag fourth-order Butterworth filter at 6 Hz) to these data. We identified heel strikes as the most anterior position of the calcaneus markers and toe-offs as the most posterior position of the second metatarsal head markers for each foot [31].

We excluded the first 45 seconds for all analyses and used the last 215 seconds, which we considered to be at “steady-state” speeds [4]. Walking speed was the sum of the treadmill belt speed and speed (0.5 Hz low-pass filtered derivative) of the approximate center-of-mass. To characterize speed fluctuations, we computed the power spectral density of the walking speeds between 0.01-0.2 Hz frequencies. Step length was calculated as the anterior-posterior distance between heel markers divided by the cosine of the treadmill angle to account for the different slopes. Step width was the mediolateral distance between heel markers for each step (heel strike to contralateral heel strike), and step frequency was the number of steps per second.

### Variability and Detrended Analysis

We computed the variances for speed, step frequency, step length, and step width. We also separated the step length and step width variances into speed-dependent (speed-trend) and speed-independent (detrended) components [26]. For each self-paced condition, we fitted the speed data with Eq. 2, to calculate the speed-trend step length based on Grieve & Gear, 1966.

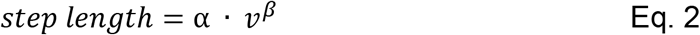

where *v* is walking speed and *α* and *β* are constants, and fitted the speed data with Eq. 3, to calculate the speed-trend step width based on Collins and Kuo, 2013.

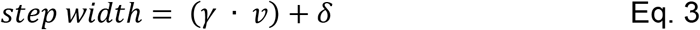

where *v* is walking speed and *γ* and *δ* are constants. No speed-related trend has been reported or is expected for step width, whereas step length has a well-established relationship with speed and is expected. For each fixed speed condition, we combined data from the three self-paced sensitivities for a slope before performing the fits to identify a single set of parameters (*α, β* and *γ, δ*). We calculated the speed-trended step lengths and speed-trended step widths using the fits (Eq. 2, Eq. 3) and subtracted them from the actual step lengths and step widths, respectively, to obtain the detrended component [26]. Then, the variance of the speed-trended and detrended components were calculated. The sum of the variance of both components should approximate to the total variance of the actual step length and step width because the speed-trended and detrended values are uncorrelated.

### Statistics

To test our main hypothesis and determine if controller sensitivity was a main effect within a given slope, we applied general linear models with repeated-measures independently for each spatiotemporal gait parameter. The assumptions of a repeated measures ANOVA (normality, homoscedasticity, and sphericity) were validated using a Shapiro-Wilk Test, Bartlett’s test, and Mauchly’s test. Outliers based on the default methods (Hi leverage, Cook’s distance, and DFITS) used in Minitab (version 19.2020.1, Minitab, LLC, State College, Pennsylvania) were excluded. Because gait experiments typically use a single sensitivity, we applied the same statistical approach with each sensitivity to determine if slope (decline, level, and incline) was a main effect for a given sensitivity. Whenever sensitivity or slope was a main effect, we used post-hoc pair-wise Tukey’s Honest Significant Difference (HSD) tests adjusted for multiple comparisons (Dunn-Bonferroni correction) to identify which sensitivities were significantly different within a given slope and which slopes were significantly different for a given sensitivity. We set the significance level to 0.05 and only report Tukey HSD p-values when either sensitivity or slope was a main effect for their respective statistical models. Additionally, we used a Wilcoxon signed-rank test to compare the fixed-speed modality and the low, medium, and high sensitivity controllers. We used SPSS (version 25, IBM Corporation, Armonk, NY, USA) for Mauchly’s sphericity test and Wilcoxon signed-rank test, and Minitab for all other statistical analyses.

## Results

### Speed Fluctuations

In all slopes, self-paced walking sensitivities had evident speed fluctuations over time, unlike fixed speed walking (Fig. 2A). Speed fluctuations were distributed across more frequencies and had greater power at higher frequencies as self-paced controller sensitivities increased within each slope (Fig. 2B). The spectral powers for the fixed speed walking (not plotted in Fig. 2B) were 2 orders of magnitude less than self-paced walking.

**Figure 2:**
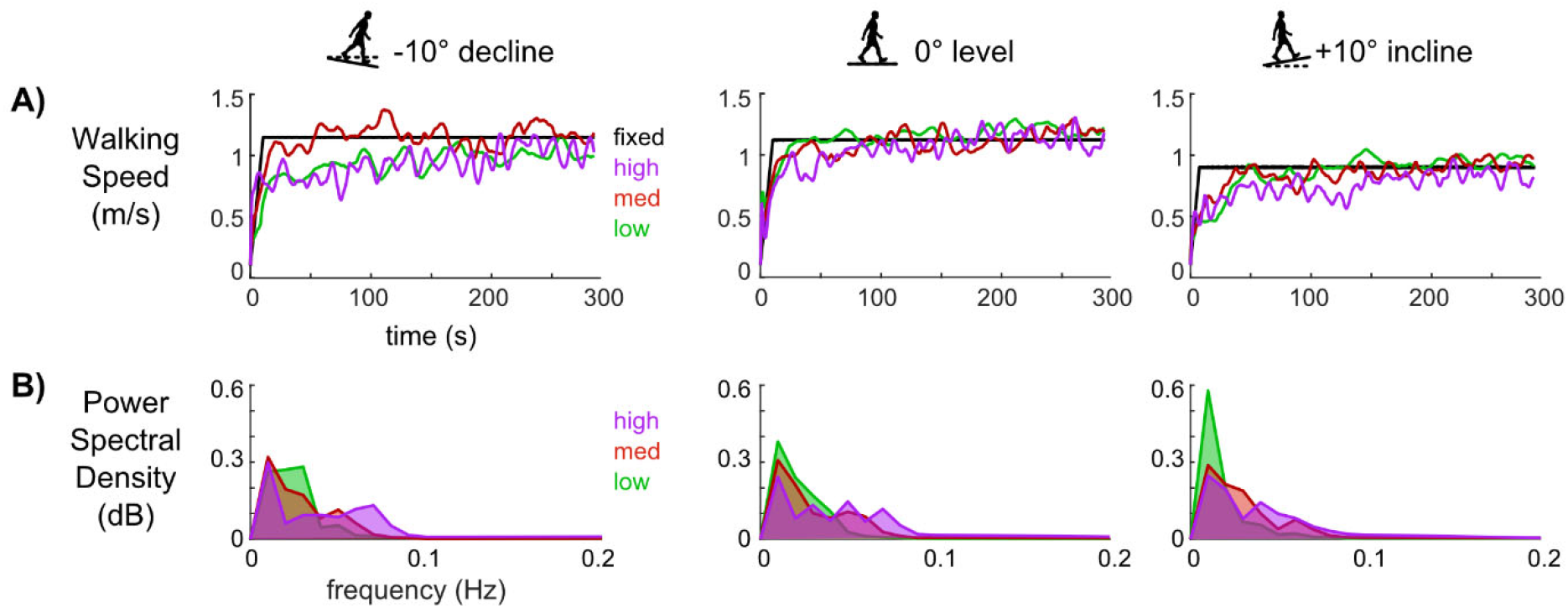
Walking speed fluctuations for a representative subject and group-average power spectral densities of walking speed. **A)** Walking speed time courses for self-paced walking with the low (green), medium (red), and high (purple) sensitivity controllers and for fixed-speed walking (black) at each slope for a representative subject. Fixed speed walking had nearly no evident fluctuations compared to self-paced walking. **B)** The group (n = 10) averaged power spectral density of walking speed was distributed across more frequencies for self-paced walking with higher sensitivities.

### Average Spatiotemporal Gait Parameters

For the level slope, the walking speeds (mean±standard deviation) were 1.24±0.28 m/s, 1.23±0.26 m/s, and 1.22±0.28 m/s for low, medium, and high sensitivities, respectively, which were similar to the 10-meter walk speeds, 1.27±0.11 m/s. Within each slope, increasing self-paced controller sensitivity did not affect average speed, step frequency, step length, or step width (Fig. 3).

**Figure 3:**
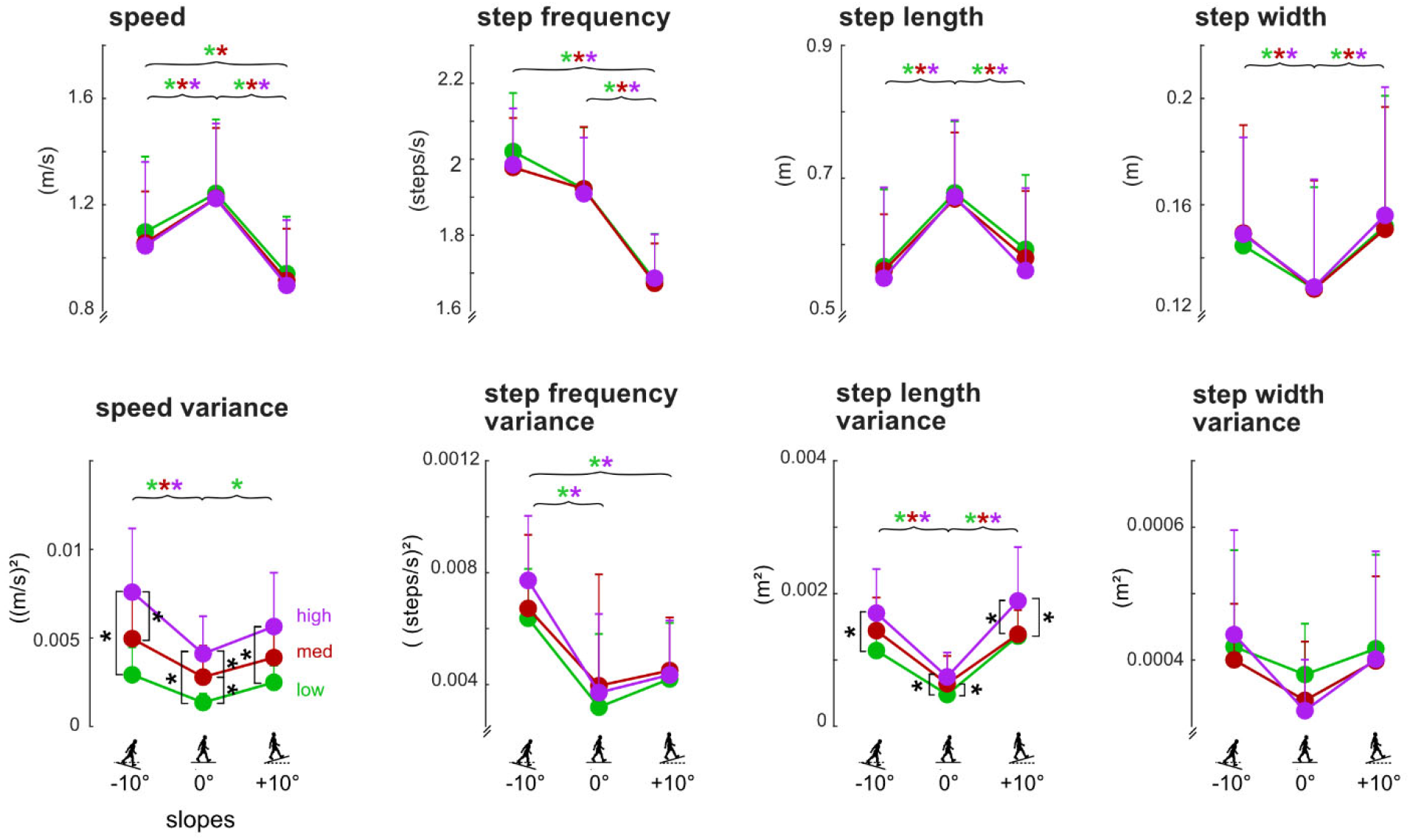
Group averaged walking speed, step frequency, step length, and step width (top row) and corresponding variances (bottom row) for each controller sensitivity for each slope. For all gait parameters, n=10 except for speed variance and step length variance where n=9, due to statistical outliers. Single sided error bars are shown and indicate standard deviations. Square brackets with black asterisks indicate significant differences between sensitivities within a slope (Tukey HSD, p<0.05). The brackets in these plots show that controller sensitivity only resulted in significant increases for speed variance and step length variance. Curly braces indicate significant differences between slopes within a single sensitivity indicated by the color-coded asterisks (Tukey HSD, p<0.05). The curly braces in these plots show that **A)** walking speed was fastest and speed variance was lowest on the level slope; **B)** step frequency was slowest on the incline while step frequency variance was highest on the decline; **C)** step lengths were longest and had the least variance on the level slope; and **D)** step width was smallest on the level slope while step width variances were not significantly different among slopes.

Each sensitivity showed that subjects walked the fastest with the longest step lengths and narrowest step widths on the level slope compared to the decline and incline slopes (Fig. 3, colored asterisks, p’s<0.05). Additionally, the slowest step frequencies were on the incline compared to level and decline slopes (p’s<0.05). With the low or medium sensitivity, subjects walked faster on the decline compared to the incline (p’s<0.05).

### Variability of Spatiotemporal Gait Parameters

Within each slope, increasing self-paced controller sensitivity did not affect step frequency variance and step width variance (Fig. 3). Only speed variance and step length variance showed significant increases with higher sensitivities within each slope (Fig. 3, black asterisks, p’s<0.05).

Each sensitivity showed that step length variance was smallest on the level slope compared to the incline and decline slopes (Fig. 3, colored asterisks, p’s<0.05), and there were no significant differences in step width variance among slopes (Fig. 3). Each sensitivity also showed that speed variance on the decline was larger than level (p’s<0.05). Speed variance on the incline was larger than level with just the low sensitivity. With the low or high sensitivity, step frequency variance on the decline slope was larger than level and incline slopes (p’s<0.05).

### Detrended Analysis

In a representative subject, greater fluctuations in speed and step lengths were evident as self-paced controller sensitivity increased (Fig. 4A), and as expected, there were strong relationships between step length and speed during self-paced walking for all sensitivities (Fig. 4B). For fixed speed walking, there was not a relationship between step length and speed, as speed was nearly constant and step lengths were concentrated in narrow range (Fig. 4A & 4B). Thus, the speed-trend step length variance for fixed speed walking was negligible (Fig. 4C). For this subject, the speed-trend step length variance was smallest for the low sensitivity controller while the medium and high sensitivities had larger but similar speed-trend step length variances (Fig. 4C). Detrended step lengths, which equaled the speed-trend step length minus the actual step length, had similar variances across the self-paced and fixed speed conditions (Fig. 4D).

**Figure 4:**
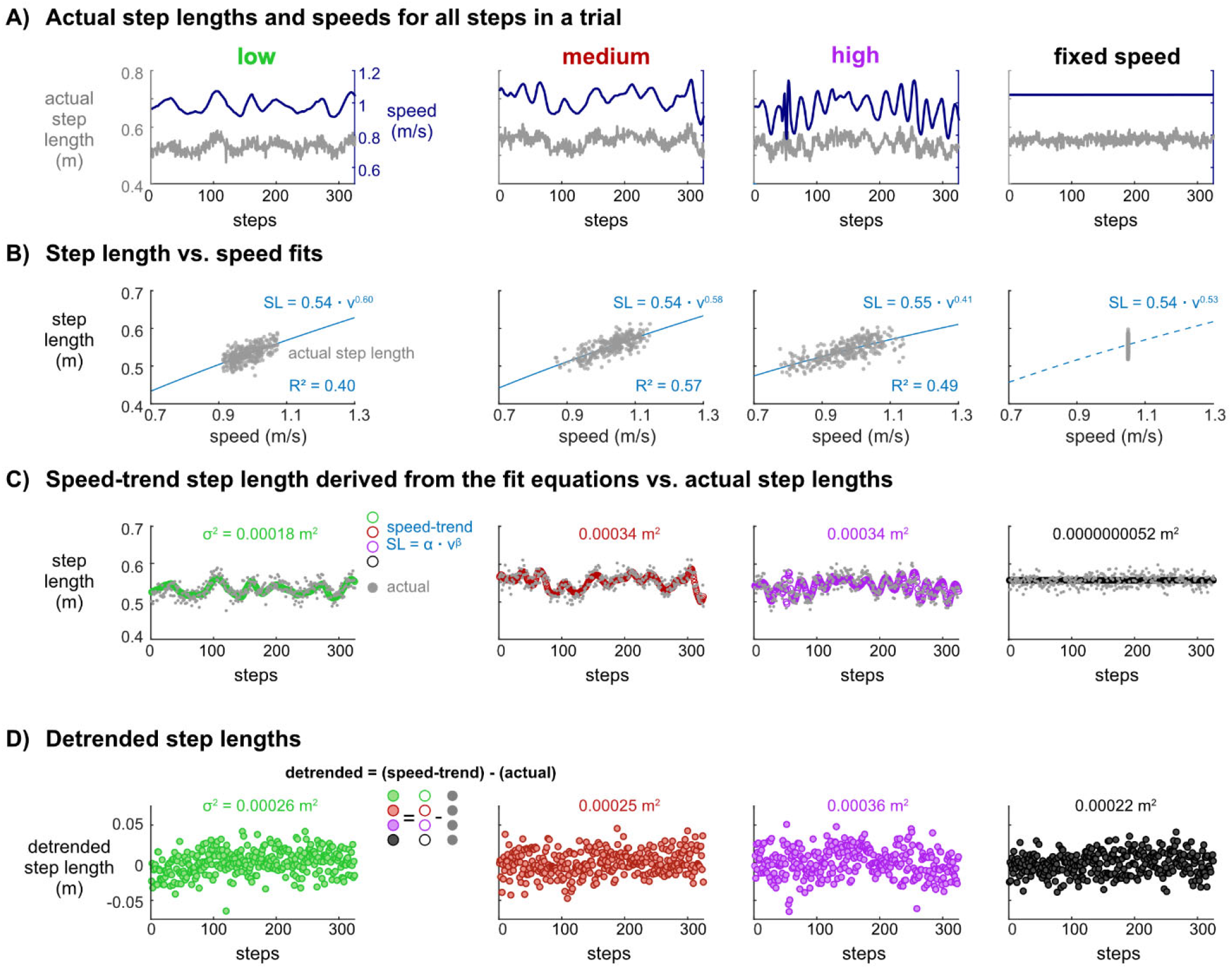
Progression of detrending speed from step length to obtain detrended and speed-trend variabilities for walking with low (green), medium (red), and high (purple) self-paced controller sensitivities and at a fixed speed (black), for a representative subject. **A)** Actual step lengths and speeds for all steps in a trial. Time courses of actual step lengths (gray lines) and walking speed (dark blue lines) paralleled each other. **B)** Step length vs. speed fits. Fit lines of the expected step length and speed relationship (Eq. 2) during self-paced walking (blue line) plotted with the actual step lengths (gray circles). The equation and line shown for the fixed speed data is from the average of the equation constants of the low, medium, and high sensitivity fit equations. **C)** Speed-trend step length derived from the fit equations vs. actual step lengths. The speed-trend step lengths (open circles) calculated from the fit equations and the actual step lengths (gray circles) are plotted for all steps. The variance, *σ*^2^, of the speed-trend step lengths was nearly zero for fixed speed walking (5.2×10^−9^ m^2^) and ranged between 0.00018 m^2^ - 0.00034 m^2^ for the self-paced controllers. **D)** Detrended step lengths. The detrended step lengths (shaded circles) are equal to the speed-trend step lengths (open circles) minus the actual step lengths (gray circles). The variance, *σ*^2^, of the detrended step lengths ranged between 0.00022 m^2^ 0.00036 m^2^ for self-paced and fixed speed walking conditions.

### Detrended and Speed-trend Step Length Variances

Speed-trend step length variances accounted for 34%, 47%, and 52% of the total variances for low, medium, and high sensitivities, respectively, on the decline, and similar percentages were observed on level (35%, 57%, 50%) and incline (41%, 46%, 58%) (Fig. 5A). Within each slope, detrended step length variances were not significantly different among sensitivities, while speed-trend step length variances were largest with the high sensitivity compared to the low sensitivity (Fig. 5B, black asterisks, p’s<0.05). Speed-trend step length variances were also larger with the medium sensitivity than the low sensitivity on the level slope (p<0.05).

**Figure 5:**
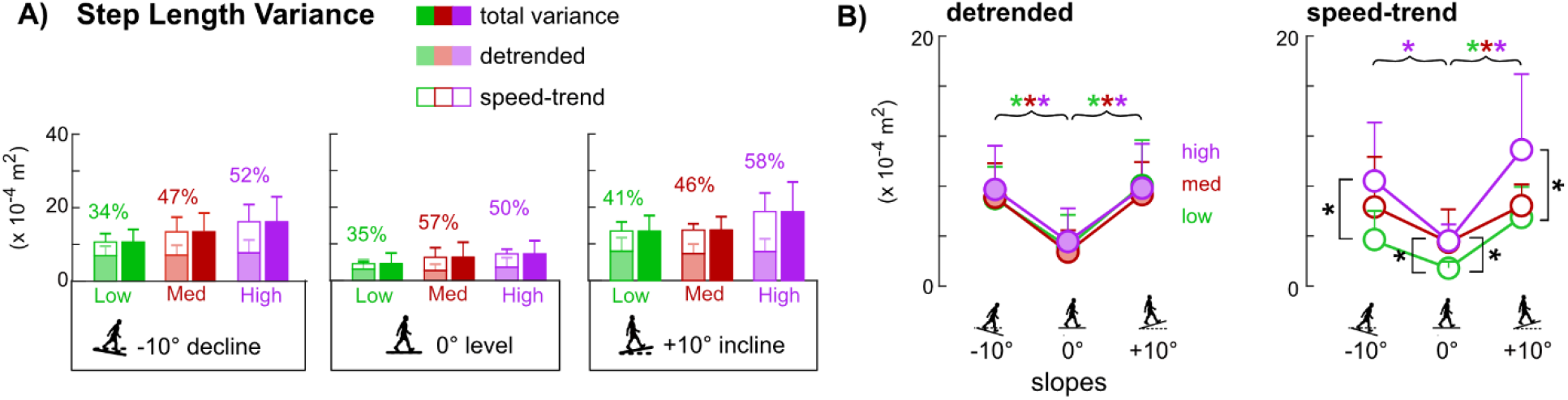
Step length variances on decline, level, and incline slopes for low, medium, and high controller sensitivities (n=9, due to statistical outliers). Error bars are standard deviations. Solid colored bars are the total variance. Stacked bars show the summation of the detrended (faded colored bar) and speed-trend (open bar) components. Percentage values above the stacked bars report the percentage of the speed-trend component. **A)** As controller sensitivity increased, the speed-trend component in total step length variance also increased within each slope. **B)** Controller sensitivity did not affect detrended step length variances within a given slope, but as controller sensitivity increased, speed-trend step length variance also increased within a given slope, as indicated with square brackets and black asterisks (Tukey HSD, p<0.05). When comparing across slopes within a given sensitivity, detrended and speed-trend step length variances were smallest on level compared to incline and decline, as indicated by the curly braces with color-coded asterisks of the specific sensitivities (green = low; red = medium; purple = high) (Tukey HSD, p<0.05).

Each sensitivity revealed that detrended step length variance was lowest on the level slope compared to decline and incline (Fig. 5B, colored asterisks, p’s<0.05) and that detrended step length variance was not significantly different between decline and incline (Fig. 5B). Each sensitivity also showed that speed-trend step length variance was larger on the incline than level (p’s<0.05). Speed-trend step length variance was larger on the decline compared to level only with the high sensitivity (p<0.05).

For the five subjects with fixed speed data, on average, fixed speed walking had smaller step length variances compared to self-paced walking (Fig. 6). Detrended step length variances for fixed speed walking were within the range of the self-paced sensitivities, while speed-trend step length variances for fixed speed walking were negligible < 1×10^−7^ m^2^ (Fig. 6). On all slopes, there were significantly greater speed-trend step length variances with the self-paced conditions compared to the fixed speed condition (p’s<0.05). However, for the detrended step length variances, there were no significant differences among the self-paced and fixed speed conditions. The total step length variances were only significantly greater with the high sensitivity controller compared to the fixed speed (p’s<0.05). However, the low sensitivity (p’s<0.10) and medium sensitivity (p’s<0.10) controllers were trending towards significantly greater total step length variance for all slopes compared to the fixed-speed condition.

**Figure 6:**
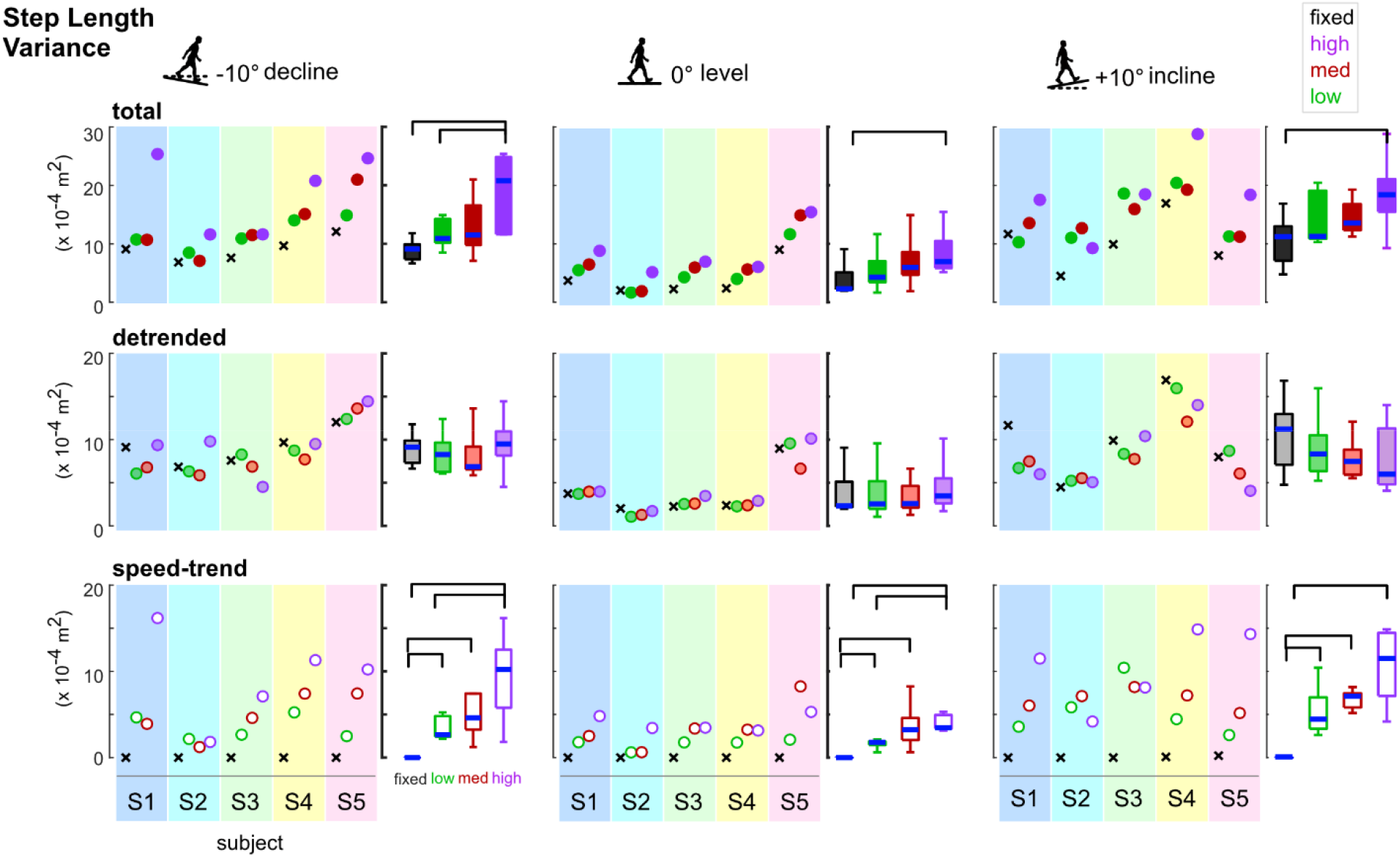
Total (solid circles), detrended (faded circles), and speed-trend (open circles) step length variances for fixed speed walking (black x’s) and self-paced walking with low (green), medium (red), and high (purple) sensitivity self-paced controllers for 5 subjects (multi-colored shaded rectangles). The box-whisker plots of the group data (n = 5) with significant differences represented by brackets (Wilcoxon signed rank test p’s<0.05). Total step length variances were significantly lower for fixed speed walking compared to the high sensitivity controller for all slopes (top row) as a result of the near zero speed-trend step length variances for fixed speeds compared to the large speed-trend step length variances with the high sensitivity controller (bottom row). Detrended step length variances were similar among fixed speed and three self-paced controller sensitivities for each slope (middle row).

### Detrended and Speed-trend Step Width Variances

Unlike speed-trend step length variances, speed-trend step width variances only accounted for 0.04%, 0.02%, and 0.05% of the total variances for low, medium, and high sensitivities, respectively, on the decline (Fig. 7A). On level and incline, speed-trend step width variances were negligible (<0.01%) in all sensitivities. Within each slope, detrended and step-trend step width variances were not significantly different among sensitivities (main effect p’s>0.05, Fig. 7B). Within each sensitivity, there were also no significant differences in detrended and speed-trend step width variances among slopes (main effect p’s>0.05, Fig. 7B).

**Figure 7:**
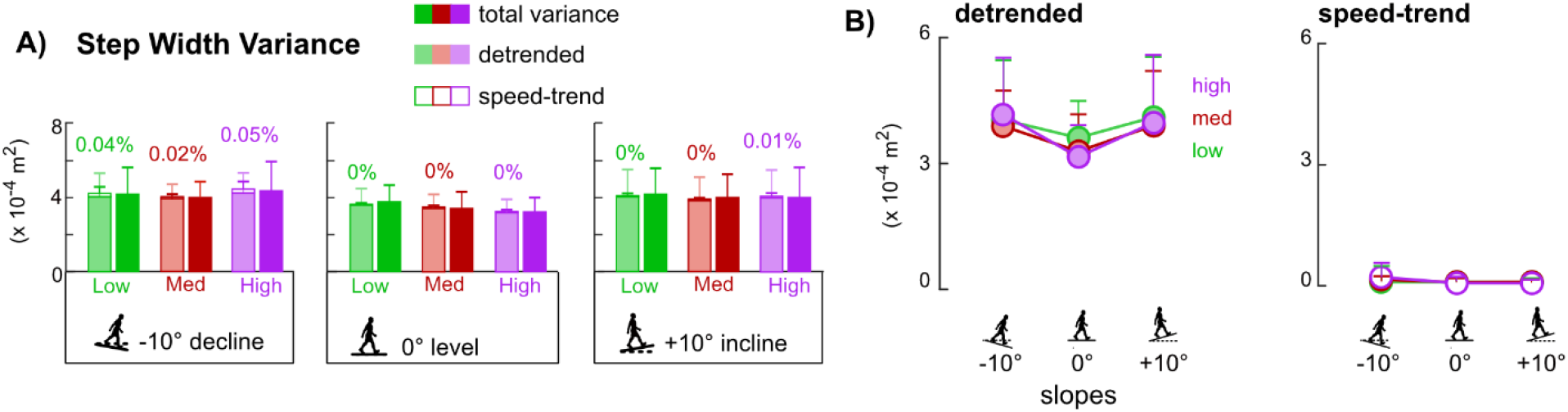
Step width variances on decline, level, and incline slopes for low, medium, and high controller sensitivities (n=9, due to statistical outliers). Error bars are standard deviations. Solid colored bars are the total variance. Stacked bars show the summation of the detrended (faded colored bar) and speed-trend (open bar) components. Percentage values above the stacked bars report the percentage of the speed-trend component. **A)** Step width variances had minimal, near zero speed-trend components within each slope. **B)** Detrended and speed-trend step width variances were not different across sensitivities within a given slope nor across slopes within a sensitivity.

## Discussion

We sought to determine how self-paced treadmill controller sensitivity affects spatiotemporal gait parameters on different slopes and to determine whether detrended variability could mitigate differences in speed fluctuations that the different sensitivities would likely impose. Within each slope, increasing self-paced controller sensitivity did not affect average walking speeds or spatiotemporal gait parameters. However, more sensitive self-paced controllers resulted in greater speed variance and step length variance but had no significant effects on step frequency variance or step width variance. Altogether, these results partially supported our main hypothesis that increasing controller sensitivity would increase speed fluctuations, step length variability, and step width variability. Importantly, as hypothesized, detrended variance was not affected by controller sensitivity suggesting that detrending variability helps to mitigate differences associated with treadmill speed fluctuations, whereas the speed-trend step length variance increased with more sensitive controllers. By detrending step length variability, the increase in total step length variance with more sensitive controllers was clearly attributed to the predominant speed-trended step length variance component. If these speed-trended contributions are not accounted for, changes in variability and potentially kinetics and energetics when walking on a self-paced treadmill may be misinterpreted. Additionally, we compared speed and gait parameters among slopes with a single sensitivity as a typical study would do. As hypothesized, walking speed was fastest and gait parameter variances were smallest on the level slope compared to decline and incline slopes with each sensitivity. Overall, our study highlights that the choice of self-paced treadmill controller parameters can alter the speed-related components of step length variability and potentially other gait parameters with strong relationships with speed. Our results suggest that detrending speed from step length variability and spatiotemporal gait parameters is one approach for accounting for the differences in speed dynamics of different self-paced treadmills.

An important finding was that self-paced treadmill walking had speed-related step length variability components that increased with more sensitive self-paced controllers for all slopes and were negligible during fixed speed treadmill walking. Differences in gait variability were not due to differences in walking speed because speed was not significantly different among the self-paced conditions in a slope and the fixed speeds matched the average self-paced walking speeds on the level slope. Multiple studies have reported that speed length variability for fixed speed walking was less than self-paced walking [5,6,12]. Our results suggest that the smaller step length variability for fixed speed walking compared to self-paced walking can be explained by the nearly non-existent speed-trend step length variance during fixed speed treadmill walking (Fig. 6). Additionally, speed-trend step length variance component and its percentage of the total step length variance increased with more sensitive self-paced controllers. Differences in gait variability would then be exacerbated with more sensitive self-paced controllers. Detrended step length variances were similar, with no systematic trends, regardless of the self-paced sensitivity or fixed speed conditions. This suggests that detrended step length variances may reveal aspects of walking control rather than being a byproduct of speed fluctuations.

Being able to interpret gait variability is beneficial for understanding walking and balance control and is one reason for striving to identify a measure that is indifferent to differences in speed fluctuations of self-paced controllers. Several self-paced treadmill studies report step variability using standard deviation, variance, or coefficients of variation but also report differing walking speeds and walking speed variabilities [4,5,7,12,21,30]. Using these measurements, step length variability was greater in self-paced walking compared to fixed speed walking, suggesting reduced balance control while potentially incurring an increased energetic cost during self-paced walking [5,12,30]. Increased variability is indeed often interpreted as being less stable [32–35] or involving greater active control [20]. Separating variability into detrended and speed-trend components, however, could additionally be interpreted as the variability attributed to steady-state walking and to changes in walking speed, respectively. Based on total step length variance or speed-trend step length variance, walking on a self-paced treadmill with higher sensitivities reduced stability and increased active control. However, based on detrended step length variance, there were no apparent differences in stability or active control between fixed speed and self-paced treadmill walking, for any sensitivity. Our results suggest that speed-trend step length variability may reflect active control needed to manage the speed fluctuations [36] and maintain the person’s position on the treadmill during self-paced treadmill walking. Detrended step length variability may reflect intrinsic active control for maintaining balance and postural control during steady-state walking, which would likely be the same regardless of treadmill features. The negligible contributions of speed-trend step width variance was expected, as lateral foot placement has little relation to speed [26]. Comparisons of step width variability across fixed speed and self-paced treadmills are likely to be less affected by any potential differences across controller sensitivities. Thus, increases in total or detrended step width variability for different treatment groups or external destabilization/stabilization manipulations on any self-paced treadmills would represent differences in mediolateral stability [35].

Our results suggest that self-paced walking on a decline was the most balance demanding while walking on level was most efficient. When walking conditions are less stable, subjects walk slowly with short and wide steps [7,37–39], which we observed on the decline and incline compared to level. The increased detrended step length variance on the decline and incline also suggests gaits were less stable compared to level because increased step length variance corresponds with greater gait instability [33,40]. Comparing between decline and incline slopes, subjects walked faster by taking more steps (i.e. increased step frequency) of similar step lengths on the decline compared to the incline. However, step frequency variance on the decline was significantly larger than incline, suggesting that decline walking was more unstable than incline walking [40–43]. Finally, self-paced treadmill walking on the level slope had the fastest speeds, longest steps, narrowest steps, and lowest variances compared to decline and incline. This could be partly due to level walking being more stable and energetically favorable compared to walking on ±10° slopes [42,44].

Similar to other studies, we found that the self-paced walking speeds matched overground walking speed [2,4]. The average difference between a subject’s self-paced treadmill speed and overground speed was ~0.04 m/s, regardless of controller sensitivity. Our average self-paced walking speed was ~1.23 m/s on the level treadmill, which is at the lower end of the range of speeds (1.23-1.61 m/s) of other studies [2,4–7,21]. Differences in speed among self-paced treadmill studies are likely due to other self-paced controller parameters instead of sensitivity, since we showed that average walking speed was indifferent to controller sensitivity on all slopes, expanding on the previous study on a level slope [6]. Differences in speeds could be related to the sample of subjects or experiment set up such as using visual optic flow. Self-paced walking speeds with optic flow was faster than without optic flow [4].

Another important consideration regarding gait variabilities across different self-paced controller algorithms is that the controller structure likely induces different speed fluctuations. Several self-paced treadmill algorithms have an overall goal of minimizing the deviations of the person’s position from an initial baseline position, which is often the center of the treadmill [1,6,18,21,36]. However, there is no standard equation for determining the treadmill belt speed corrections, and the equations vary depending on the signals related to the treadmill system that can be used in a custom written real-time controller. Published self-paced algorithms use ground reaction forces [2,21], echo location [1], or motion capture [4,6,7] to calculate the changes in a subject’s position on the treadmill. Several self-paced treadmills controllers use proportional-derivative control [6,21,36], while other controllers use proportional-integral-derivative control [18] or proportional control [45]. The frequency of the treadmill belt speed corrections varies among controllers spanning from updates at each footstep [21], at 30 Hz (for Motek’s old algorithm) [4,6], and at 120 Hz [19]. Additionally, the self-paced controller structures likely affect the magnitudes of the belt speed fluctuations and walking speed fluctuations. Several studies report the standard deviation of the self-paced walking speeds, which quantifies the spread of walking speeds, providing an indicator of the magnitude of the walking speed fluctuations. In our study, the range of walking speed standard deviations for the low, medium, and high controller sensitivities for the level condition spanned between 0.04 m/s to 0.06 m/s. In another Motek self-paced treadmill study for level conditions, walking speed standard deviations were between 0.05 m/s to 0.07 m/s [6] whereas in other studies that reported walking speed variability with custom designed self-paced treadmill controllers, the walking speed standard deviations were 0.06 m/s to 0.1 m/s [5,30] (Fig. 8).

**Figure 8.**
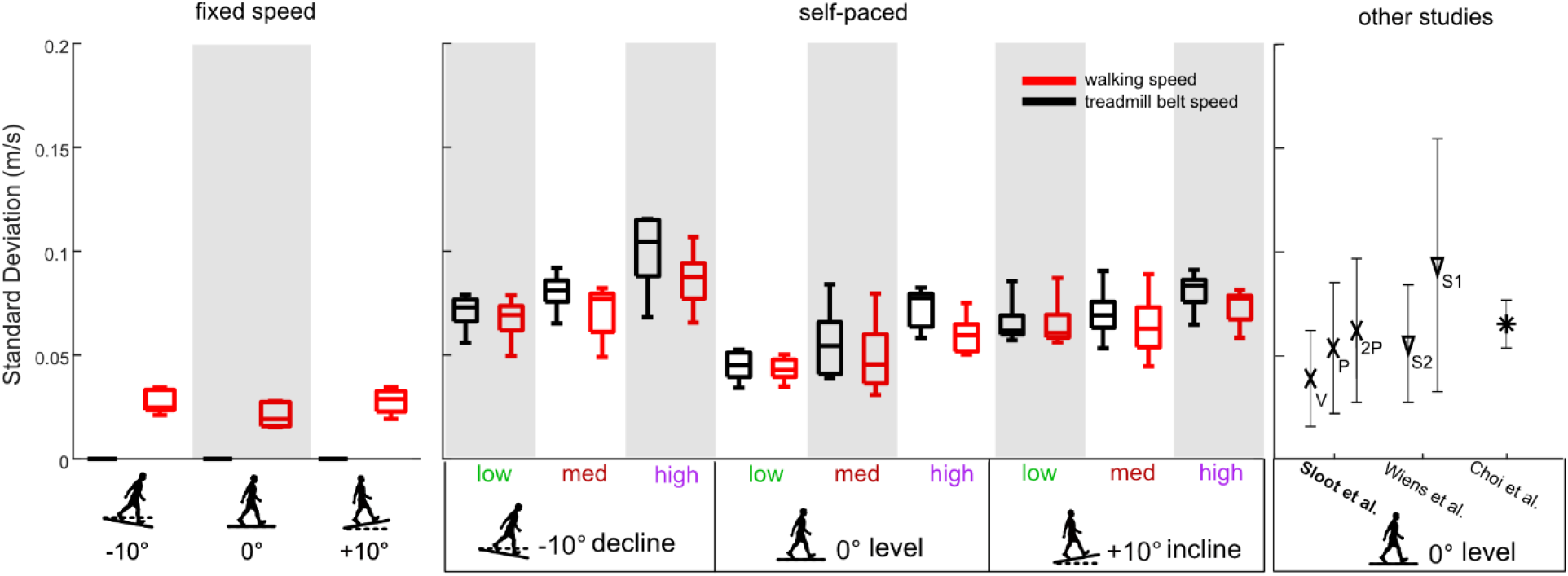
Standard deviations of walking speed (red) and treadmill belt speed (black) on decline, level, and incline slopes for our fixed speed conditions (n=5), our self-paced conditions (low, medium, and high sensitivities) (n=5), and for other studies that used a self-paced treadmill. The fixed speed conditions had treadmill belt speed standard deviations that were near zero, as expected, while walking speed standard deviations for fixed speed conditions were within the range of walking speed standard deviations for the self-paced conditions. The treadmill belt speed and walking speed fluctuations were generally similar for the self-paced conditions. Previous self-paced treadmill studies report a wide range of walking speed standard deviations. The Sloot et al. study used the old algorithm for the Motek treadmill and reported values for three self-paced controllers, V, P, and 2P, which correspond to the abbreviations in their paper [6]. The Wiens et al. study reported values for two sessions collected on separate days with the same self-paced controller, shown here as S1 and S2 [30]. Studies using the Motek treadmill are bolded.

Limitations of this study include potential learning effects of walking on a self-paced treadmill and testing a small subset of sensitivities and slopes. Participants with no experience walking on a self-paced treadmill or sloped treadmill could experience a learning effect over the nine self-paced walking conditions. Another limitation is that we only used the maximum slope angle possible, ±10° and did not test shallower slopes. Similarly, we did not test sensitivities that spanned the full range, which could have revealed additional significant effects of sensitivity. Lastly, detrended spatiotemporal gait parameters were only analyzed with linear variability measures. The applicability of detrending for nonlinear variability measures, such as detrended fluctuation analysis, would require further investigation.

In summary, self-paced treadmill walking included speed-dependent gait variability components that increased with higher sensitivity self-paced controllers. Self-paced controller sensitivity had non-significant effects on average spatiotemporal gait parameters (speed, step length, step width, and step frequency) on decline, level, and incline slopes. Detrended variability seems to mitigate potential effects of different treadmill speed dynamics of self-paced controllers on gait variability, which may be beneficial when comparing gait variability among multiple self-paced treadmill studies or with fixed speed conditions.

## Acknowledgements

We would like to thank members of the Biomechanics, Rehabilitation, and Interdisciplinary Neuroscience (BRaIN) Lab for help with data collections and discussions.

## Supplement Caption

**S1 Figure:**
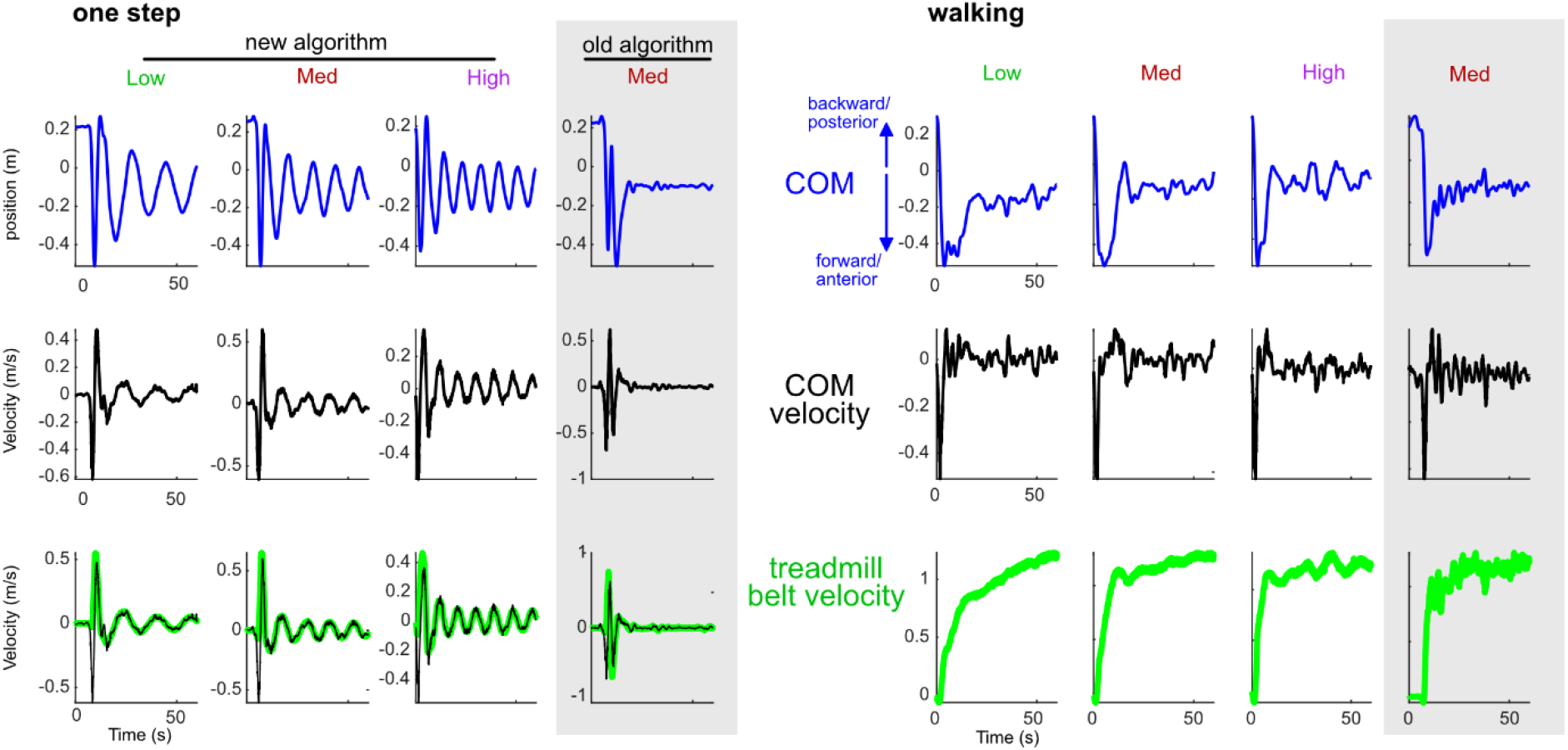
The center of mass (COM) position in the anterior-posterior direction, treadmill belt velocity, and COM velocity in the anterior-posterior direction for a single step and walking for 60 seconds using Motek’s “new algorithm” with the low, medium, and high sensitivity values used in our study and using Motek’s old algorithm (reported in Sloot et al. 2014) with the default/medium sensitivity value of 1 (gray rectangle). For the single step condition, the subject took a single forward step and then stood still on the treadmill belt. For the single step condition, after an initial correction in response to the change in the person’s relative position with the treadmill, the “new algorithm” produced treadmill belt velocity oscillations that resulted in COM position oscillations with a slow decay rate towards the steady state position whereas the “old algorithm” rapidly reached the steady state COM position. The COM oscillations increased in frequency for increasing sensitivities with the “new algorithm.” For the walking condition, the “old algorithm” had more frequent changes in the COM, treadmill belt velocity, and COM velocity compared to the “new algorithm.”

